# Germline CpG methylation signatures in the human population inferred from genetic polymorphism

**DOI:** 10.1101/2023.03.24.534151

**Authors:** Yichen Si, Hyun Min Kang, Sebastian Zöllner

## Abstract

Understanding the DNA methylation patterns in the human genome is a key step to decipher gene regulatory mechanisms and model mutation rate heterogeneity in the human genome. We analyzed existing whole genome bisulfite sequencing (WGBS) data across tissues and large genetic variation catalogs and observed that 93.2% CpGs hyper-methylated in sperm are polymorphic. Moreover, methylation status of CpGs is spatially correlated, as 94% of CpG pairs within 1kb share the same methylation status. Leveraging only these properties, we infer germline CpG methylation in the human population using a new method, Methylation Hidden Markov Model (MHMM), and the polymorphism data from TOPMed. Our inference is orthogonal to WGBS-based experimental results; still we observed 90% concordance with human sperm WGBS while overcoming several challenges in that data: We inferred methylation status for ∼ 721, 000 CpG sites that were missing from WGBS due to low coverage, and show that 42.2% of CpGs with allele frequency > 5% are hyper-methylated in the population but could not be captured in WGBS due to sample genetic variation. Our results provide a unique resource for CpG methylation levels in germline cells complementary to the existing WGBS-based measures, and can thus be leveraged to enhance analysis such as annotating regulatory and inactivated genomic regions in the germline.

## Introduction

DNA methylation can directly modify protein binding sites or change chromatin 3D organization to regulate gene expression[1], and the majority of DNA methylation in mammalian cells is contributed by CpG methylation[2]. DNA methylation is also crucial for understanding mutation processes. In the human germline, the cytosine to thymine (C>T) mutation rate at methylated CpG sites is ten fold greater on average than that of unmethylated CpG sites[3, 4], leading to the observation that 99% of methylated CpG sites are mutated in at least one of 390k individuals [5]combining genomAD[6], UK Biobank[7], and DiscovEHR[8].

Whole genome bisulfite sequencing (WGBS) is the gold standard for measuring CpG methylation level[9–11], and > 100 human tissues and cell lines have been profiled[12]. However, each of these datasets provides measurements for one sample of cells and cannot be extrapolated to population level methylation. Moreover, understanding methylation from an evolutionary perspective requires historical methylation information, which is never directly measurable. In particular, experimental data is limited by germline mutation bias[13], where a typically methylated C has mutated to a T. Individuals carrying a C>T mutation would be faithfully measured as unmethylated by bisulfite sequencing obfuscating the historical methylation at this locus or the methylation status of other individuals with a C allele. As methylated CpGs have high mutation rates, many such obfuscating mutations have reached high allele frequencies in the population. Across 45 million autosomal CpG sites across the genome, random individuals are expected to carry 805,979 C>T mutations and to have homozygous T alleles at 135,222 sites, based on allele frequencies from Bravo[14]. This mutation bias reduces estimates of the mean methylation level, especially in small samples.

WGBS is especially challenged when estimating germline methylation. Germline methylation is crucial to understand developmental processes and germline mutations. Germline methylation pattern is best estimated from sperms, oocytes, and germline cells at early developmental stages[4, 15]. Among them, methylation status in sperm has the strongest correlation with germline mutation rate and SNP density in population samples[4]. However, although researchers have reported high methylation rate and distinct methylation patterns in sperm[4, 9], the number of available germline WGBS data is too small (1 sperm, 2 testis, and 3 ovary samples published on ENCODE[16], all from different studies) to allow conclusive statements about human germ cell methylation[15, 17]. Because DNA methylation is dynamic during development and differs across tissues[15, 17, 18], combining information from WGBS datasets across tissues or cell lines will not mitigate the difficulties in studying germline methylation.

On the other hand, computational approaches have been developed to identify genomic features that affect CpG methylation level and predict DNA methylation[19–22]. For example, Zhang et al.[20] use a variety of genome annotation, especially histone marks and regulatory elements, to train a statistical random forest predictor for methylation level in whole blood, for which epigenetic experiments have the largest sample sizes over multiple modalities. DeepCpG[22] combines DNA sequence context and incomplete methylation measures to impute missing methylation status using neural network in single cell data. Both methods borrow information from observed methylation in a neighborhood to infer missing methylation status at a focal CpG position.

We propose a new method to infer germline methylation level independent from experimental methylation measures, using observed allele frequencies in publicly available variant catalogs at single base resolution. Allele frequency is informative about germline methylation status at a CpG site as methylated sites have very high mutation rates. As a result of their ∼ 10 fold increase of mutation rate, some CpG sites that are consistently methylated in the population have mutated multiple times (recurrently) in the sample history, so that hyper-methylated regions are depleted of monomorphic sites and low-frequency variants (Figure 1a). Along the DNA sequence, both methylated and unmethylated sites tend to form tight regions with high or low methylation rates so that information can usefully be shared locally (Figure 1b). For instance, CpG islands, empirically defined as CpG dense segments, are highly enriched in genic region and protein binding sites and are often non-methylated. In contrast, methylation of consecutive CpGs in a promoter is a mechanism to silence the corresponding genes[23, 24]. Combining this information, we developed Methylation Hidden Markov Model (MHMM), which infers hidden germline methylation levels at individual CpGs sites from allele frequencies of C>T variants (Figure 1c).

**Figure 1:**
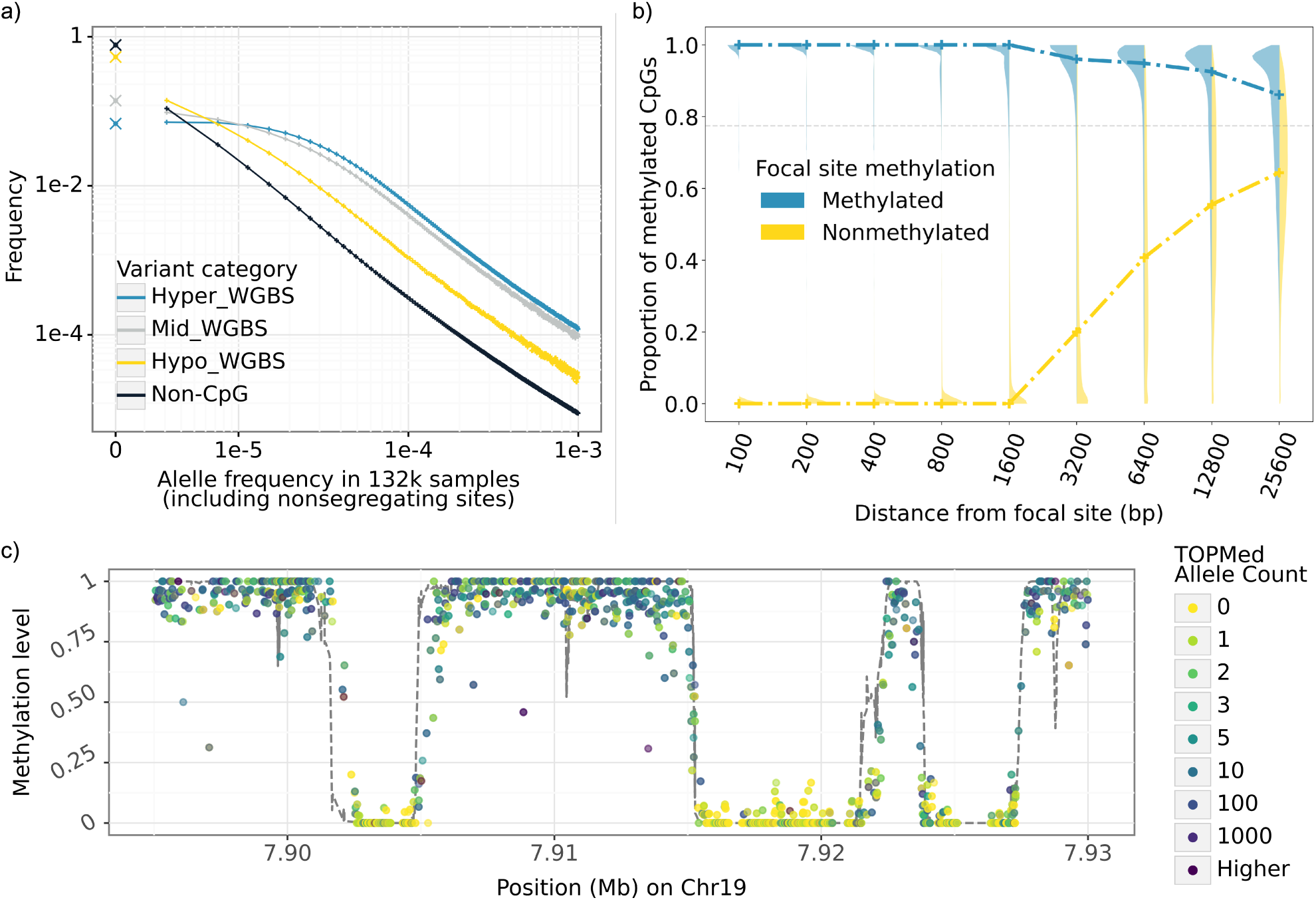
Data patterns of CpGs leveraged for inference in MHMM. a) Difference of sample frequency spectrum (SFS) between hyper-methylated (blue), hypo-methylated (yellow), and intermediate (gray) CpGs in a sample of 132k individuals informs the emission probabilities of the HMM. SFS among non-CpG sites (black) is provided for comparison. Crosses mark the fraction of monomorphic sites. Methylation status is based on WGBS measured sperm methylation level. b) Proportion of hypermethylated CpGs around a focal CpG site as a function of the distance from the focal site informs the transition probabilities of HMM. Violin plots show the distribution of this proportion in varying neighborhood sizes; blue and yellow dashed lines mark the median values when the focal site is hyper- or hypo-methylated respectively. Within a small distance of a hyper-methylated (hypo-methylated) CpG, most CpGs are hyper-methylated (hypo-methylated); beyond 20kb from a hypo-methylated CpG the average methylation rate is close to the global average (gray horizontal line). c) An example 40kb region on chromosome 19 showing the raw data. X-axis is the genomic position in Mb, each point is one cytosine in a CpG site colored by its allele count among 132k individuals from the TOPMed study[14]. Y-axis for the points is the methylation level measured by WGBS in sperm. Dashed line is the MHMM inferred probability of being hyper-methylation using only the allele counts as observations.

We apply MHMM to whole genome polymorphism data on 132,345 individuals from the TOPMed study[14] to infer population level germline methylation. Although our model uses information orthogonal to experimental measures or sample specific methylation status, our results are consistent with sperm methylation level measured by WGBS at 90% of CpG sites and our inferred hypo-methylation CpGs are highly enriched in known active genomic regions. Since our results can also be interpreted as accumulated mutation burden at near base pair resolution, contrasting the observed and expected allele frequencies suggests CpG sites that are likely to be under purifying selection. Our software and inferred methylation levels are available at https://github.com/Yichen-Si/cpghmm.

## Results

### Overview of the experiments

We apply our Methylation Hidden Markov Model (MHMM) to infer germline methylation levels in humans using the TOPMed variant catalog[14] (freeze 8, 132,345 genomes) and replicate our observations in gnomAD[6] (v3.0, 71,702 genomes). We compute the probability distribution over discretized methylation levels at each of the 45 million autosomal CpG loci conditional on all observed CpG allele frequencies (AF) within 20Mb or to the end of the chromosome arm.

Our estimates are highly consistent with the methylation status measured in sperm cells by WGBS, while differences between MHMM and WGBS indicate both limitations of our method and blind spots in WGBS. We demonstrate potential applications of the method, showing that 1) Inferred hypo-methylated CpGs are enriched for active/regulatory genomic regions; 2) CpG sites located in inferred hyper-methylated regions but monomorphic in the sample are enriched for sites where C>T mutations would cause severe functional consequences (Figure 2).

**Figure 2:**
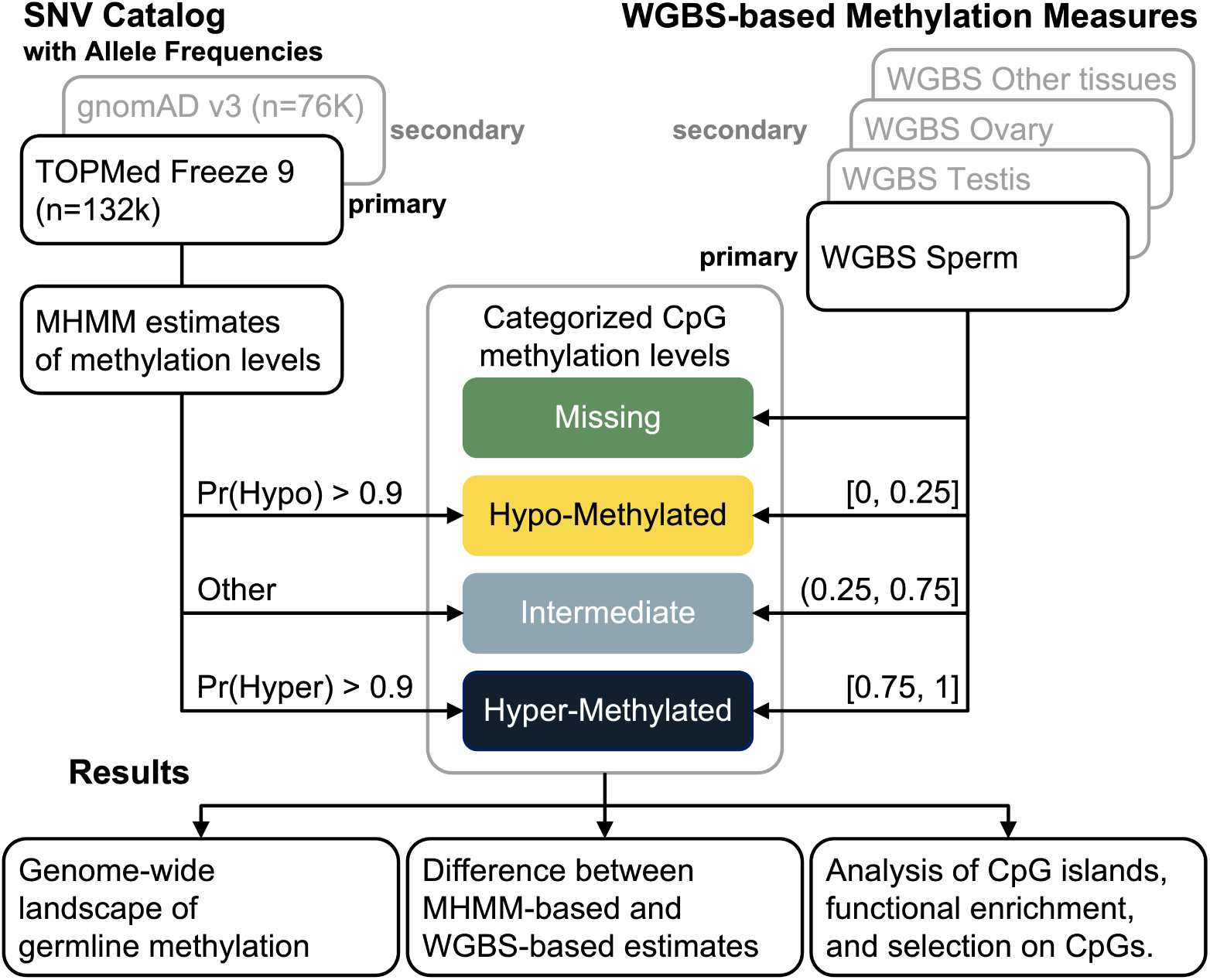
Overview of the analyses.

### Application of MHMM on the TOPMed variant catalog

When applied to the TOPMed variant catalog, MHMM assigns most CpG sites to one of the two categories: 77.7% of CpG sites have a > 0.9 probability of being in hyper-methylation regions, 12.0% of CpG sites have > 0.9 probability of being in hypo-methylation regions while the remaining 10.3% CpG sites have intermediate methylation level or cannot be confidently assigned to either category (Figure A.1). The inferred methylation level is associated with local CpG densities, with hypo-methylated regions enriched in dense CpG neighborhoods (Figure A.1, A.2). The difference between maximum likelihood (MLE, using all observed CpG AF) and leave-one-out likelihood (LOO, excluding the focal site’s own AF from its estimator) is generally small, exceeding 0.1 (probability unit) at only 6.4% of sites. This difference is on average higher in regions with lower CpG density, ranging from 0.025 to 0.018 across density deciles.

### Replication of MHMM results with gnomAD

We applied the same model fitting process to gnomAD[6] AF based on 71,702 individuals primarily of European and African ancestries. Results from gnomAD are consistent with those from TOPMed in general (Table A.1) with 90.6% CpGs having the same categorical methylation levels inferred from the two variant catalogs. There are 0.73% CpGs inferred as hyper-methylated using one dataset while inferred as hypo-methylated using the other, the other 8.72% discrepancy is the result of one of the datasets suggesting an intermediate methylation level. The general consistency is expected since AF do not differ qualitatively between the two variant catalogs except for rare variants. But the two datasets differ in sample sizes and in rare variant calling and quality control procedures, both affecting the lower end of the site frequency spectrum (SFS) where most signal for our model is from. Henceforth, we present results from TOPMed which has a larger sample size.

### Relationship between CpG methylation and SFS is tissue-specific

The high mutation burden of CpGs consistently methylated in the germline result in a distorted SFS. Figure 1a shows the SFS of CpGs stratified by their methylation status in sperm measured by the whole genome bisulfite sequencing (WGBS), compared with SFS of non-CpGs. Only 6.8% of hyper-methylated CpGs remain monomorphic among 132k individuals (TOPMed freeze 8 from Bravo[14]), distinct from hypo-methylated CpGs where 53.6% remain monomorphic. Figure 3 shows that the SFS of hyper-methylated CpGs differs by the tissue where the methylation status is measured. CpGs hyper-methylated in sperm are most enriched for polymorphic sites, followed by those hyper-methylated in testis tissues. Ovary tissue shows similar level of enrichment to other non-germline tissues. This observation is consistent with previous estimates that male contributes 4 times germline mutations than females[25, 26]. We also observe that tissue samples show more intermediate methylation levels, 18.1 ∼21.3% among 5 samples of testis or ovary compared to 7.7% in the sperm cell line sample. This observation is consistent with the fact that these two tissues commonly used as proxies for germline cells[5] consist of multiple cell types. These observations demonstrate the limitation of using experimental data from tissue samples including testis and ovary to understand germline methylation. Therefore, we compare MHMM inferred methylation levels with the WGBS measure of a sperm sample as the best approximation of germline methylation unless stated otherwise. In this sperm dataset, among 55.3M autosome CpGs 69.4% are hyper-methylated (with measured methylation rate ≥ 0.75), 14.8% are hypo-methylated (with measured methylation rate ≤ 0.25), and 8.1% are missing (Figure A.3).

**Figure 3:**
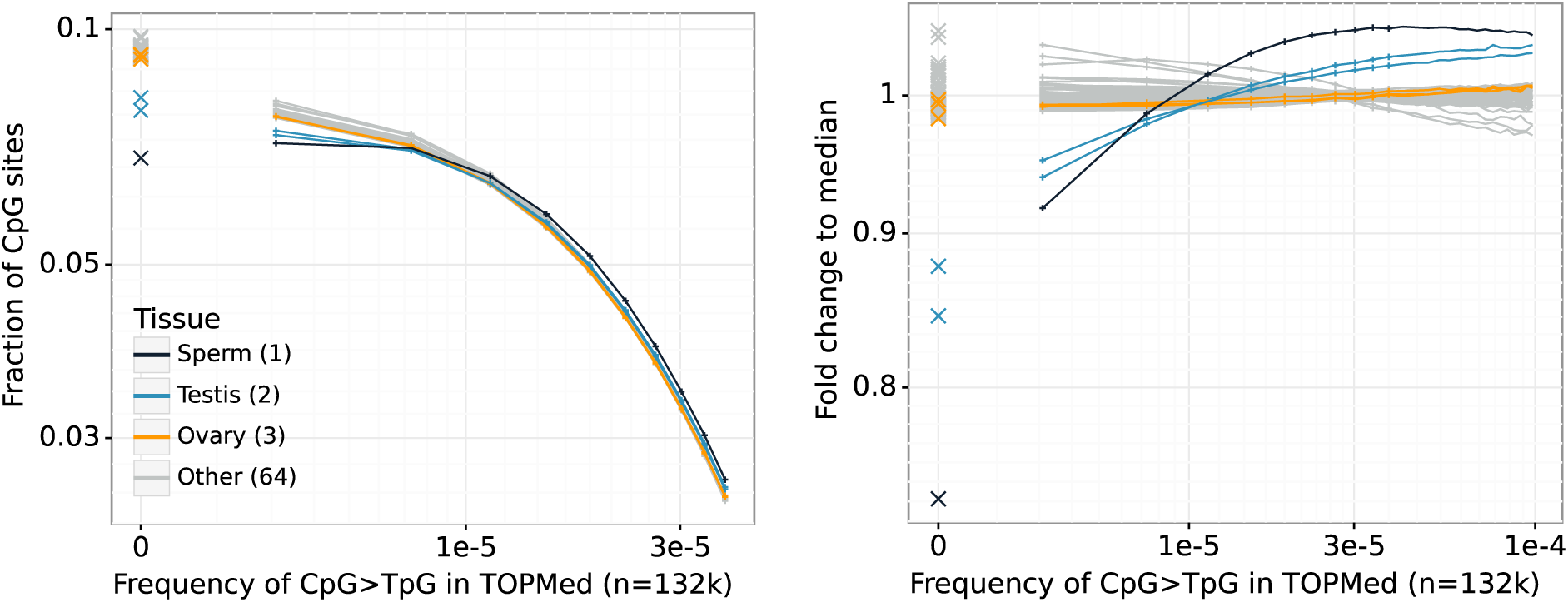
SFS of CpGs hyper-methylated in different tissues. Left: SFS truncated to highlight the rare variant tail (x- and y-axis are in log scale); right: fold difference of the SFS compared with the median among non-germline tissues (x-axis is in log scale). Each line is one sample, those from sperm, testis, and ovary are colored as black, blue, and orange; gray lines are non-germline tissues. The left most points in both figures represent monomorphic sites in Bravo.

### Comparison between MHMM inferred and WGBS measured methylation levels

Comparing inferred germline methylation with WGBS measured sperm methylation, we see that among inferred hyper-methylated CpGs, 90.0% are measured as hyper-methylated by WGBS and 1.7% are measured as hypo-methylated by WGBS; among inferred hypo-methylated CpGs 90.1% are measured as hypo-methylated and 3.6% are measured as hyper-methylated. Among the remaining 10.7% CpGs inferred as having intermediate methylation level, 59.8%, 19.6%, and 18.4% are measured as hyper-methylated, intermediate, and hypo-methylated respectively (Figure 4a).

**Figure 4:**
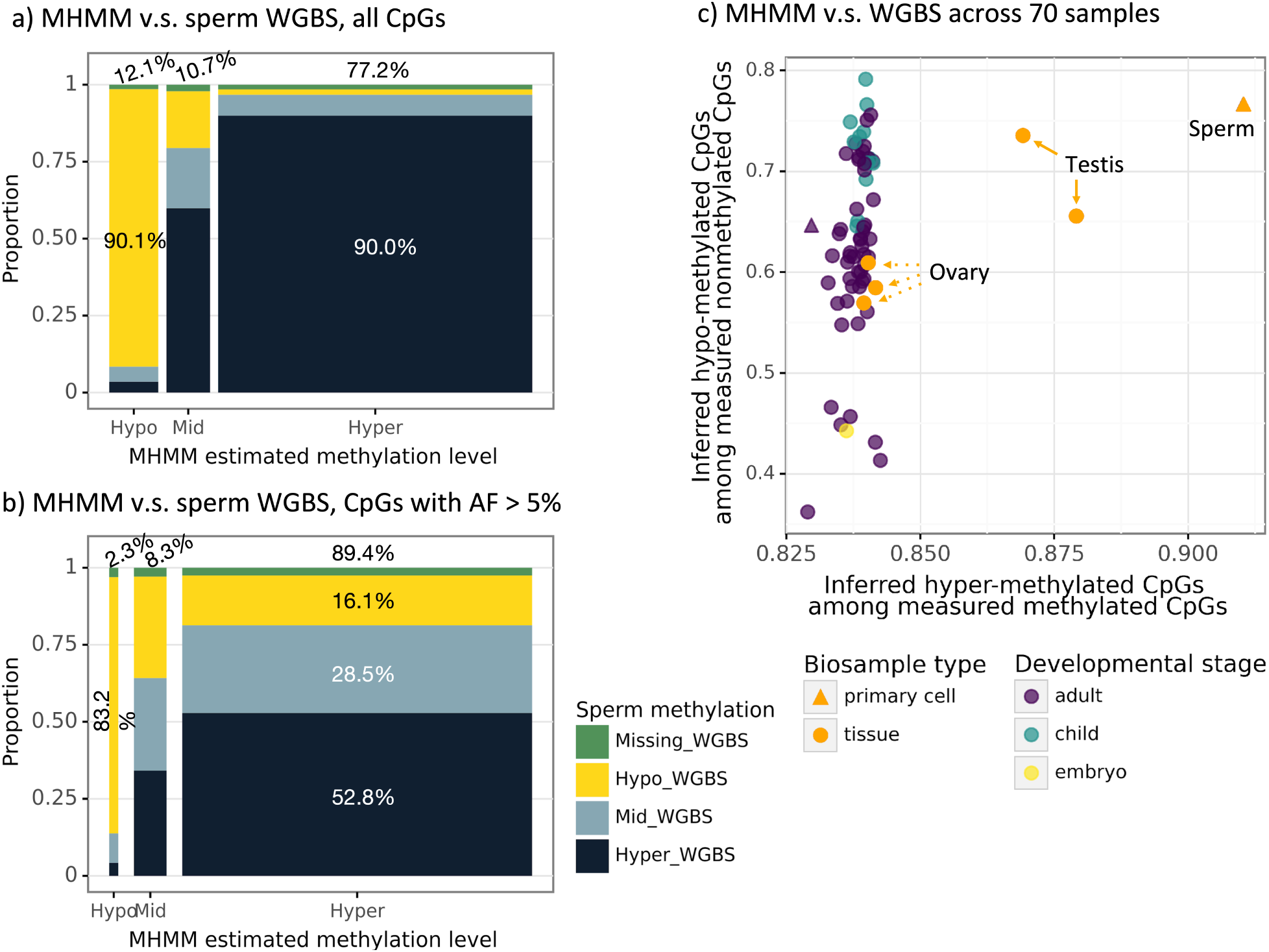
a) Compare MHMM inferred methylation level with WGBS measures in sperm across all autosome CpGs. Width of each bar is proportional to the number of CpGs in the corresponding MHMM inferred methylation category, i.e. the denominator of the proportion on the y-axis. b) is similar to a) but includes only CpGs that are common variants with AF> 5%. c) Comparison between MHMM estimated methylation level and that measured by WGBS across 70 samples. Each point is one sample; colors indicate the ages of donors of non-germline samples, and germline samples are highlighted by orange (all from adults). All but two samples are tissue samples, the sperm data we focus on is a primary cell sample (triangle).

We extend this comparison to other 69 samples from diverse tissues and cells (Table A.4) and observe that among those other samples, the testis tissues’ methylation levels are the most similar with MHMM inference, although less so than sperm. Ovary tissues’ methylation levels are less similar with MHMM inference, in fact they are comparable with other non-germline tissues (Figure 4c).

To better understand the 10% discordance between MHMM and WGBS, we identify three sequence properties associated with such differences. First, discordance depends on local CpG density. In the 10% sparsest regions (less than 10 CpG per 1kb), 74% of measured hypomethylated sites are inferred to be intermediate or hyper-methylated. In contrast, in the 10% densest regions (≥60 CpG per 1kb), 6% of measured hypo-methylated sites are inferred to be intermediate or hyper-methylated. This difference in concordance is likely driven by the fact that the MHMM integrates information across neighboring CpGs but the correlation of methylation levels decays rapidly with distance[20, 22] (Figure 1b). As GC content is higher in coding sequence and near transcription start sites (TSS) compared to intronic and intergenic regions[24], our inference and WGBS agree more in genic regions (Figure A.4). Second, WGBS missing values are distributed unevenly across the genome. After removing regions with low mappability or low sequencing quality (see Method), 1.6% of the remaining 45.2M CpG sites have missing methylation status, and CpG sites with higher local GC content or higher AF are enriched for missing values. Among 5.86M CpGs located in 65,551 autosomal CpG islands[27] 7.7% are missing WGBS observations (4.8x enrichment), and among 176,257 CpGs with AF >0.5, 3.2% are missing WGBS observations (2.0x enrichment).

### Germline mutation bias affects WGBS-based methylation estimates

Third, The discordance between MHMM and WGBS is also enriched among CpGs where the mutant T alleles have high frequencies (Figure 4b). Because experimental techniques correctly read T alleles at mutated CpGs as non-methylated, inference of population level methylation based on a small sample of individuals is biased by their germline mutations[13]. Here we assess this bias also in the WGBS sperm sample to disentangle this mutation effect from differential methylation between tissues or cell types.

We categorize CpG sites by their T-allele frequencies from Bravo[14] and show the distribution of WGBS methylation level in each AF window (Figure 5). Among monomorphic (AF = 0) sites and ultra-rare variants (AF < 0.01%) 17% are hypo-methylated while among intermediate frequency variants (AF 0.01% ∼ 1%) 4% are unmethylated. This is consistent with the fact that hypo-methylated sites have lower mutation rates and so are less likely to be polymorphic. However, among common variants (AF >1%) 31% of CpGs measured as partially methylated or unmethylated, and 57% of high AF variants (AF >50%) are measured as hypo-methylated. This contradicts what is expected based on mutation rate but is consistent with the donor often carrying the mutant T alleles at high AF variant sites.

**Figure 5:**
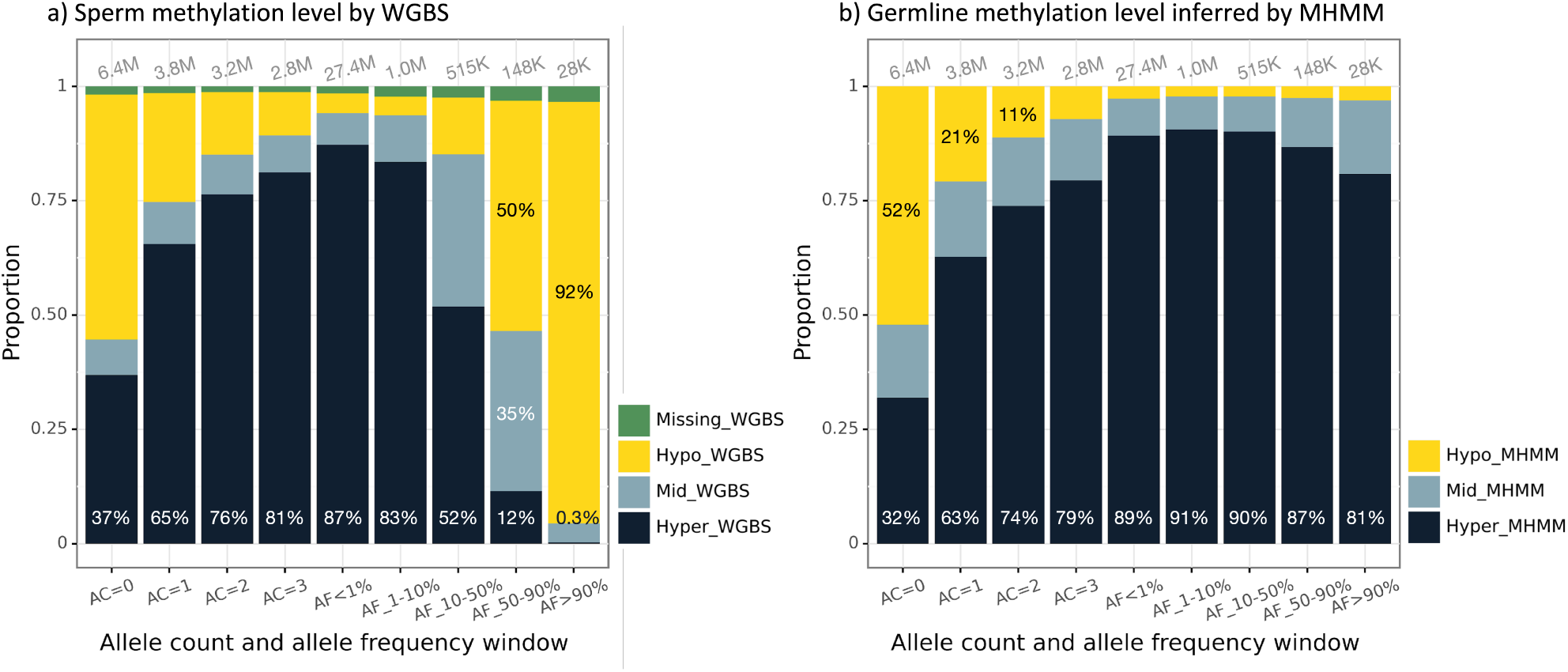
WGBS measured (left) and MHMM inferred (right) germline methylation level distribution by AF. We divide whole autosome reference CpG, including monomorphic sites, into 10 (unequal sized) bins from rare to common (x-axis), and show the proportion of methylation levels according to either WGBS or MHMM.

Further, while methylation status is generally similar among nearby CpGs, CpGs with high T-allele frequencies measured as hypo-methylated by WGBS are typically surrounded by hyper-methylated CpGs. For each CpG site we assess the methylation status of its 5 immediate CpG neighbors both upstream and downstream as measured by WGBS. Among hypo-methylated focal sites with AF <0.01, only 8.0% have any of their 10 neighbors hyper-methylated. In contrast, among hypo-methylated focal sites with AF >0.9, ≥95.3% of sites have at least one of their 10 neighbors hyper-methylated. For 61.0% of such sites, all of their 10 neighbor CpGs are hyper-methylated (Figure A.5), suggesting that these sites would likely to be methylated if the C alleles are intact, but are measured to be hypo-methylated only because the sample donor carries the T allele.

The germline bias leads to biased SFS stratified by CpG methylation levels. CpG methylation is commonly used as a predictor for germline mutation rate, which is in turn an important factor to account for when estimating selection signal at gene or region level[6]. Figure A.6 highlights the enrichment of intermediate and high frequency variants among CpGs measured as hypo- and mid-methylated by WGBS due to the germline bias. Since germline hypo-methylation usually indicates low mutation rate and low expected variant density, the enrichment of common variants deviates from the expectation and may create false signal for balancing or positive selection.

This germline bias in WGBS contributes to ∼10% of the discordance between MHMM inferred germline methylation level and the observed methylation level. Across all CpGs 1.7% of MHMM inferred hyper-methylated CpGs are measured to be hypo-methylated by WGBS; but among inferred hyper-methylated CpGs with AF 10-50% and AF>90%, 9.1% and 93.6% are measured as hypo-methylated by WGBS. In addition, 37.2% of inferred hyper-methylated CpGs with AF 10-90% show intermediate methylation level by WGBS, compared to only 7.7% of estimated hyper-methylated CpGs with AF below 10%, consistent with the large proportion of heterozygous genotypes in the AF range (Figure A.7). If we project the proportion of hyper-methylated CpGs among sites with intermediate AF to those with high AF, we expect ∼367,000 CpG sites hyper-methylated in the population to be mutated in the donor and therefore their methylation status obfuscated.

### MHMM-based methylation levels separate known CpG islands into two classes

CpG islands are defined as regions that have high local GC content and enriched CG dinucleotides compared to the expectation if nucleotides are distributed randomly[27]. Although 70% of annotated promoters in the human genome are located in such high CpG density regions[28], whether a CpG island can function as transcription regulator heavily depends on its methylation status[24]. We observe that known CpG islands fall into two distinct groups characterized by their inferred germline methylation level, consistent with previous studies evaluating sequence conservation[29]. Among 25,743 autosome CpG islands, 77.1% are primarily (>90%) hypo-methylated while 13.4% are primarily (>90%) hyper-methylated. Among hypo-methylated CpG islands 42.2% overlap with known promoters and 37.0% overlap with known proximal enhancers (Table A.2). Among hyper-methylated CpG islands, only 0.3% overlap with known promoters and 0.6% overlap with known proximal enhancers.

### Enrichment of hypo-methylated CpGs in regulatory regions

We further assessed the functional annotations of all hypo-methylated CpGs identified by either or both of MHMM and WGBS. MHMM and WGBS identify 5.4M and 6.4M hypo-methylated CpGs respectively, where 56% MHMM-hypo and 62% WGBS-hypo CpGs are located outside known CpG islands. We calculated the enrichment (odds ratios (OR)) of hypo-methylated CpGs in regulatory and active genomic elements, including TSS (MHMM OR=35.7), proximal enhancers (OR=3.6), open chromatin (OR=43.1), and transcription factor (TF) binding sites (OR=12.1) (Figure 6). CpGs identified as hypo-methylated by MHMM show slightly higher enrichment than those identified by WGBS and CpGs identified as hypo-methylated by both MHMM and WGBS show slightly higher enrichment than that identified by either method alone.

**Figure 6:**
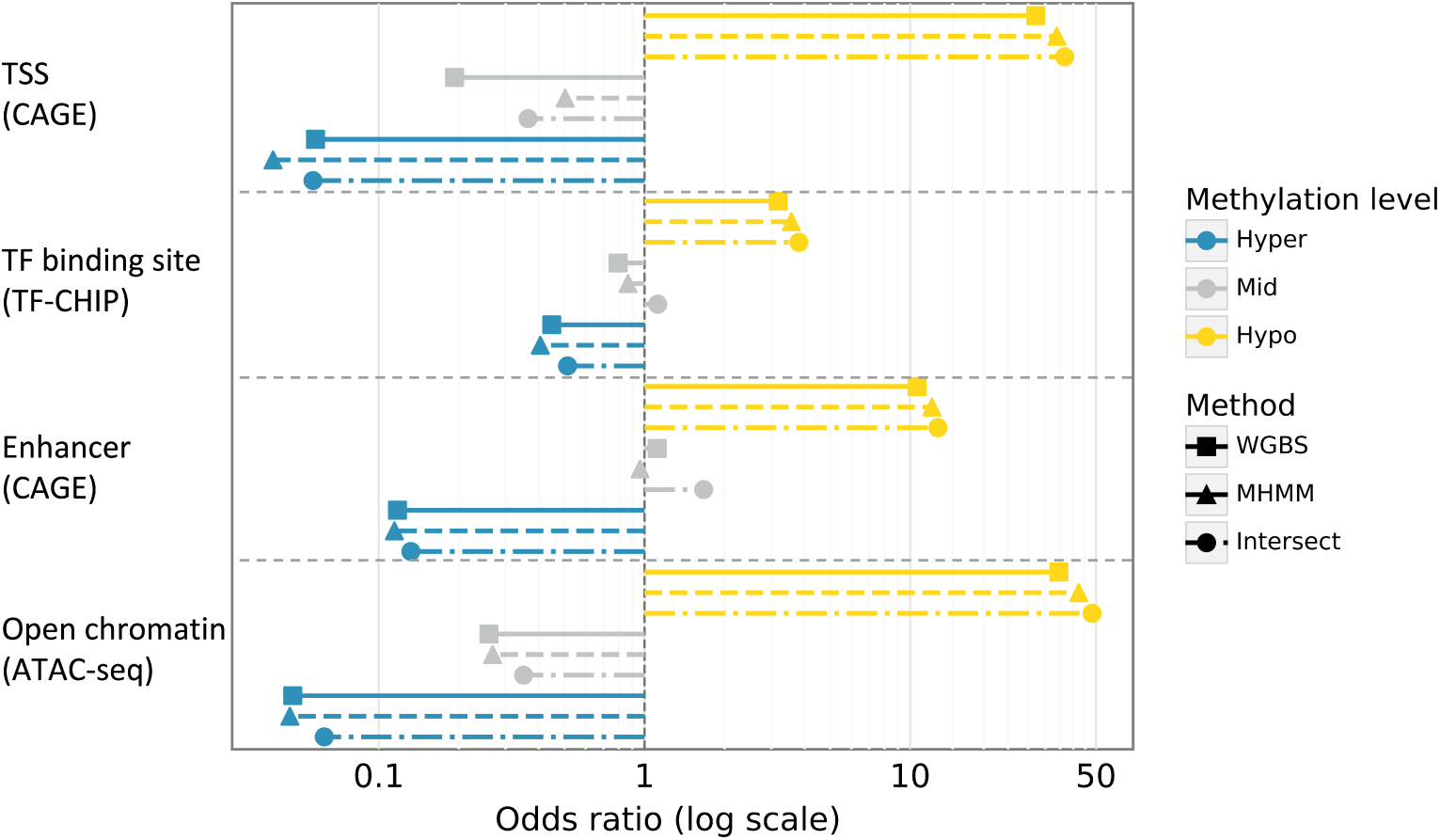
Enrichment of known active and regulatory elements among hypo-methylated CpGs. X-axis is the odds ratio of the enrichment in log scale, from top to bottom shows the enrichment of TSS, enhancers, TF binding sites, and open chromatin (data sources see B). Rectangles and triangles show the odds ratio when the methylation status is defined by by WGBS and MHMM respectively. Circles show the odds ratio with the subset of CpGs where WGBS and MHMM methylation status are the same. (The approximated confidence intervals are negligible in relation of the scale of the odds ratios)

### Monomorphic hyper-methylated CpGs show strong signatures of purifying selection

The segregation pattern of genetic variations in a population sample is shaped not only by the mutation rate, but also by purifying selection. We classify CpG sites by the potential effect of C>T mutations using the Ensembl Variant Effect Predictor (VEP)[30] and stratify SFS among hyper-methylated CpGs by the functional consequences of the C>T mutation (Figure 7). Among all hyper-methylated CpGs 94.5% are polymorphic and 26.6% have an AF ≥0.01%. Among putative loss of function (LoF) mutations, 71.6% sites are polymorphic and only 7.0% reach AF ≥0.01%, suggesting strong purifying selection. We used the leave-one-out estimator of MHMM (see Method) to estimate the methylation level of such highly selected CpGs without the constraint on allele frequency confounding the estimate (Figure A.8).

**Figure 7:**
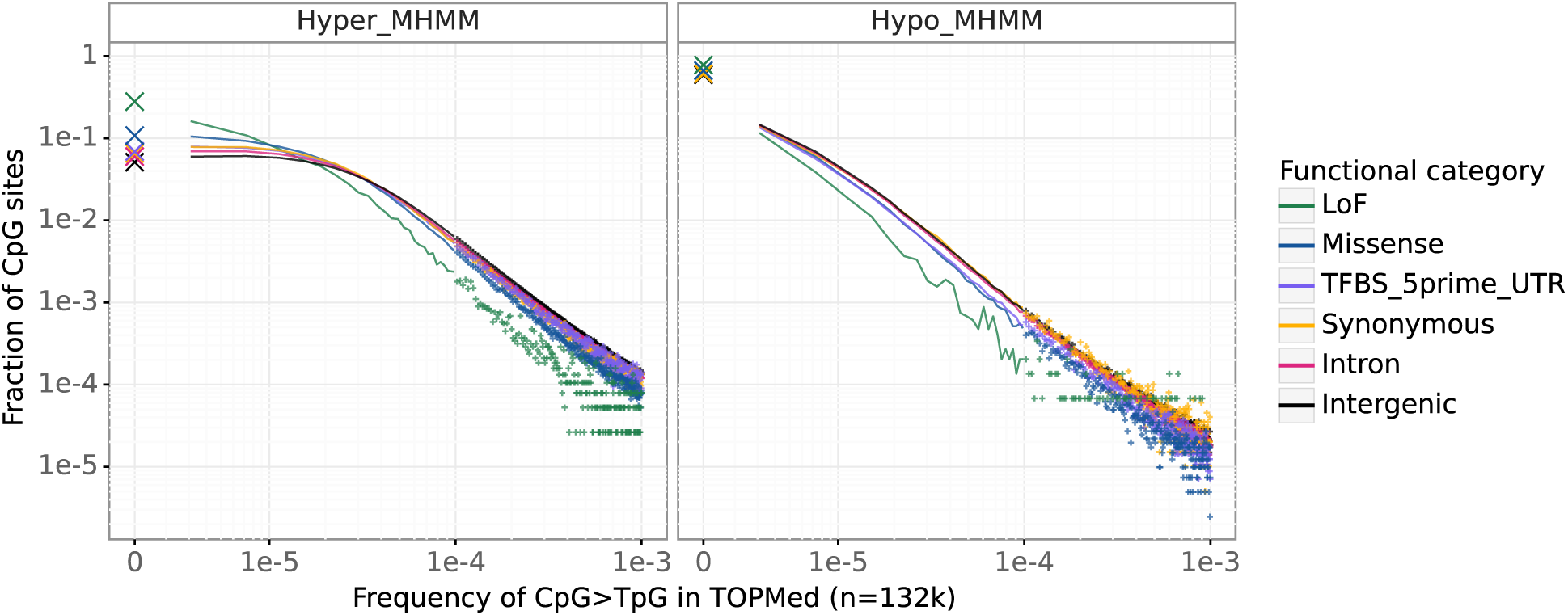
SFS of CpG C>T mutations stratified by MHMM inferred methylation levels and predicted variant effects (from Ensembl VEP[30]). Left: hyper-methylated CpGs, right: hypo-methylated CpGs. Frequency of monomorphic sites are marked separately by crosses.

This difference of SFS by predicted variant function suggests that hyper-methylated but monomorphic CpGs are likely to be under purifying selection. Table A.3 shows the proportion of monomorphic site by mutation consequences and methylation levels. Among CpG sites where a C>T mutation would be synonymous, 6.6% of hyper-methylated CpGs remain monomorphic, compared to 61.0% among hypo-methylated CpG sites, a > 9-fold difference due to the latter’s low mutation rates. Among CpG sites where a C>T mutation is predicted to cause the loss of function (LoF) of the protein, 27.9% of hyper-methylated CpGs remain monomorphic, compared to 78.1% among hypo-methylated CpG sites, < 3-fold difference because purifying selection partially cancels the mutation rate effect.

## Discussion

We developed a Methylation Hidden Markov Model (MHMM) to infer population level germline methylation at CpG dinucleotides. Our method leverages the high mutation rates and the consequently distorted sample frequency spectrum (SFS) at hyper-methylated cytosines and the local correlation of methylation status. We applied MHMM to polymorphism data from the TOPMed database and inferred whole autosome CpG methylation. Among CpGs inferred to be germline hyper-methylated by MHMM 90% are measured as hyper-methylated by WGBS in sperm cells, and 93.2% of CpGs hyper-methylated in sperm are polymorphic in the TOPMed.

The differences between inference by MHMM and experimental measures by WGBS are driven both by limitations of MHMM and blind spots in WGBS. As MHMM aggregates the signal from nearby CpGs, it has limited information for inferring the methylation level of isolated CpGs. Indeed, MHMM differs from WBGS at about 74% hypo-methylated CpGs in the 10% lowest density regions but only at 5% hypo-methylated CpGs in the 10% highest density regions. On the other hand, WGBS’s precision is limited in two scenarios. First, originally hyper-methylated CpGs that now have high derived T-alleles frequencies in a population sample may not be identified by WGBS[13]. Experimental measures from a small number of samples reflect the methylation status and the genotypes of those samples. CpG sites carrying a T allele are then identified correctly as unmethylated even if the C allele would be methylated, thus interrupting the otherwise hyper-methylated regions. On the contrary, MHMM inference reflects the population level methylation signature, is independent of individuals’ genotypes, and is more locally homogeneous.

Second, bisulfite sequencing often has low coverage in GC-rich regions and thus low quality methylation calls. This GC content bias is more significant with PCR amplification which is currently standard in WGBS[11]. In the sperm data (GSM1127119) for example, we have observed a 4.8 fold enrichment of missing data in known CpG islands compared to the global missing rate. The polymorphism data underlying our methylation inference is primarily from PCR-free sequencing protocols therefore are less affected by GC content bias.

Thus, MHMM creates more comprehensive estimates of germline methylation especially in highly mutable and GC-dense regions. and combining MHMM inference with the experimental data improves our ability to identify germline open chromatin and regulatory elements that are often hypo-methylated. MHMM identified hypo-methylated CpGs are more likely to locate in known active genomic regions than WGBS-identified hypo-methylated CpGs, and hypo-methylated CpGs supported by both methods are more likely to locate in known active genomic regions than those supported by either method alone. Since MHMM is orthogonal to WGBS experiments in the source of both information and bias, one can further explore alternative strategies to integrate the two methods based on local GC density and allele frequency.

To the best of our knowledge, MHMM is the first method that infers methylation independent of experimental assays. All exiting prediction methods are supervised methods that rely on and thus share the limitations of the technologies like WGBS. Comparing our method with other computational methods is also difficult because these methods typically use tissue or cell type specific genomic features unavailable for germline cells. For example, Zhang *et al.* (2015) achieved 91.9% prediction accuracy of methylation level in blood cells, which have the largest sample size and most complete genomic annotations, by using various functional features including each focal site’s neighboring methylation status.

Since SFS is shaped by mutations happened in the past generations in the sample genealogy, MHMM estimates can be interpreted as the methylation signature accumulated in the population history. The strong deviation of SFS from the expected SFS under the low, genome-wide averaged mutation rate is a signature of CpGs consistently methylated over generations thus maintaining high germline mutation burden in the population. The high consistency of the current day sperm methylation level in a random individual with this accumulated signature inferred by MHMM also suggests that human germline methylation pattern is relatively stable in the past and across the population.

Based on this evolutionary interpretation, we show that some CpG sites are maintained by purifying selection[5]. A CpG site where its own allele frequency substantially differs from that predicted by its neighborhood is 3.7 times more likely to be a potential loss of function mutation site. To account for purifying selection in our inference, we calculate two statistics, the marginal and leave-one-out likelihoods, for methylation status at single base pair resolution. We adopted a heuristic combination of the two statistics that effectively reduces confounding from purifying selection while incurring minimal loss of information at the majority of CpG sites that are near neutral.

Although we emphasize the population level interpretation of the inferred methylation signature, a future application is to compare our results with more individual measures from human and closely related species. For instance, our inferred germline methylation level could serve as a baseline to be compared with that measured in specific tissues to identify patterns acquired during developmental processes; or with that measured in a small sample from a distinct population to investigate germline variation across ancestries; or with that measured in chimpanzees, gorillas, or orangutans to detect local fast evolution.

When applying our method to study germline methylation patterns in different human groups or non-human species, high quality sequencing data are critical. An intrinsic limitation of our method is its reliance on accurate rare variant genotyping to capture the number of monomorphic sites and ultra-rare alleles, because this lower tail of the SFS is the most informative about germline mutation rate. To be robust to sample demographics and genotyping artifacts, we do not constrain the SFS by population genetics parameters, but instead let the model learn the difference of SFS across genomic regions from the data.

Overall, we demonstrate that we can leverage the accumulated mutation burden at CpG sites to infer germline methylation level averaged over past generations and across the human population, and reveal methylation patterns hidden from experimental measures. Our results also provide a new resource for interpreting non-coding regions by identifying potential regulatory elements and CpGs under mutation constraint.

## Availability

Our inferred methylation levels using Bravo and genomAD respectively, and a track for UCSC Genome Browser are available at https://zenodo.org/records/10140747. Our software is available at https://github.com/Yichen-Si/cpghmm.

## Acknowledgement

We thank Dr. Eleanor J. Clowney for discussion on the biology of methylation and CpG islands.

## Author contributions

Y.S., H.M.K., and S.Z. developed the ideas; Y.S. and S.Z. wrote the manuscript with contribution from H.M.K.; Y.S. implemented the software, and performed the analyses.

## Method

### Hidden Markov Model

To estimate the unobserved methylation status from polymorphism data, we build a continuous time, discrete state HMM. We model discretized methylation levels as hidden states, discretized allele frequencies as observations, and construct the transition probability as a function of the physical distance between adjacent CpG sites. The stochastic process consists of all individual cytosines in CpG sites on both strands of a chromosome according to human reference genome GRCh38, and the emission models local sample frequency spectrum (SFS) as a result of the local methylation level.

We discretize methylation level into *K*(≥ 2) states from hypo- to hyper-methylated, and the T-allele frequencies into *M* categories from monomorphically C to primarily T. Let *Z_i_* and *X_i_* indicate the hidden state (methylation level) and observation (allele frequencies) at location *i* respectively, so *Z_i_ ∈* [*K*] = {1*, . . ., K}*, *X_i_ ∈* [*M*]. We explicitly include monomorphic (allele count AC= 0), singleton (AC= 1), doubletons (AC= 2), up to AC= 5 as separate categories, then choose log-linearly spaced break points to group higher allele frequencies. In practice, the two tails of the sample frequency spectrum (SFS) are most informative for our inference. We choose *K >* 2 because the historical methylation level averaged over time and population is likely to be fractional rather than binary as in a homogeneous cell sample. Choosing a state space with finer resolution also increases the robustness to mutation rate variation among hyper-methylated CpGs that depends on sequence context and other unmodeled factors[31].

We assume that the emission probability of a single observation conditional on its state follows a multinomial distribution, *P* (*X_i_* = *m|Z_i_* = *k*):= *E_k,m_*, where *E_k,·_* corresponding to a binned SFS specific for state *k*. We do not assume any structure of *E* and its estimates are dataset specific because methylation-level-stratified SFS depends on population demographics and sample size.

We parameterize transitions among states in two layers. At each position (the C in a CpG), we first generate its probability of moving out of the current state by the next position based on an exponential distribution of the distance until a change of state, then sample the next state if a transition occurs. We consider two adjacent CpG sites *i* and *i* + 1 separated by *d_i_* base pairs (bp). The probability for position *i* + 1 to stay in the same state as position *i* conditional on *z_i_* = *k* is

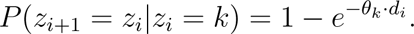

Conditional on *z_i_*_+1_ ≠ *z_i_* the probability to move to state *k′* ≠ *k* is *A_k,k′_*, where *A* is a stochastic matrix satisfying Σ_*k*≠*k*_ *A_k,k′_* = 1*, A_k,k_* = 0 for all *k, k^′^ ∈* [*K*]. Taken together, the transition from *k* to *k*^′^ ≠ *k* over *d_i_* bp has probability

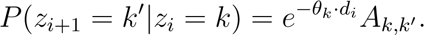

We assume the *K* transition rates *θ_k_*are independent, capturing the different length distribution of hyper- and hypo-methylation regions; we do not constrain conditional probabilities *A* to any structure, allowing the relation among states to be learnt entirely from the data.

### Parameter estimation

Unknown parameters in this model include *A, E,* and *{θ_k_}*, with *K*^2^ *−* 2*K* + *KM* degrees of freedom. Matrices *A* and *E* can be estimated with the Baum–Welch algorithm[32], but the maximum likelihood estimates for transition rate *θ_k_*’s do not have analytical forms. We use an approximate EM approach and update *θ_k_*’s iteratively by numerical optimization. We choose the Nelder–Mead method, a gradient-free, multi-dimensional optimization algorithm to search for a local optimum in the K-dimensional parameter space of *θ_k_*’s conditional on *A* and *E* in the M-step of each iteration. We estimate the parameters by running the above EM algorithm on each arm of the autosomes separately (excluding centromeres), then take the genome wide average of learned parameters weighted by the number of CpG sites in each chromosome arm.

Modeling observed allele frequencies as the sole result of underlying methylation level is subject to confounding from natural selection. Monomorphic CpG sites in a large sample could have low methylation levels and low mutation rates, or could be under purifying selection that removes C>T mutations[33]. To reduce the effect of selection on the inference of individual CpG sites, we compute a leave-one-out (LOO) likelihood of the hidden states at each CpG conditional on all but the focal site’s allele frequency. In this way, we avoid assuming neutrality at every single site and instead assume it is rare to have a dense cluster of CpG sites all under selection that dominate a neighborhood. This assumption is mild because even in highly selected coding regions, fitness effect of CpG C>T mutations is limited by amino acid changes possible in the context. Computing LOO likelihoods only adds a fraction of overhead using similar computational tricks as in[34, 35].

### Data application

We train the above model and estimate marginal and LOO likelihoods for each CpG cytosine to be at each of the discretized methylation levels using autosomal allele frequencies from Bravo (132,345 genomes) and gnomAD (71,702 genomes) separately. We excluded ∼ 10M CpGs located in regions with low mappability based on the reference from the 1000 Genome Project[36] or low variant quality scores reported by the variant catalogs, resulting in ∼ 45M autosomal CpG sites.

We compare our results with methylation levels of 43 different tissues and cell lines from 80 publicly available WGBS datasets from the ENCODE portal[16] (https://www.encodeproject.org/, individual dataset identifiers are listed in supplementary). To best approximate the historical methylation we aim to estimate, we focus on a sperm cell line from a 28yr male (ENCSR705FPH) which is the only sperm sample from ENCODE[16]. We did not include female germline because the ratio between male and female contribution to germline mutations is about 4:1[26] and the available proxy for female germline, ovary tissue samples, contain mixtures of cell types with high non-germline contribution. We annotated the functional consequences of C>T mutations at CpG sites using Ensembl VEP[30].

## A Supplementary figures and tables

**Figure A.1:**
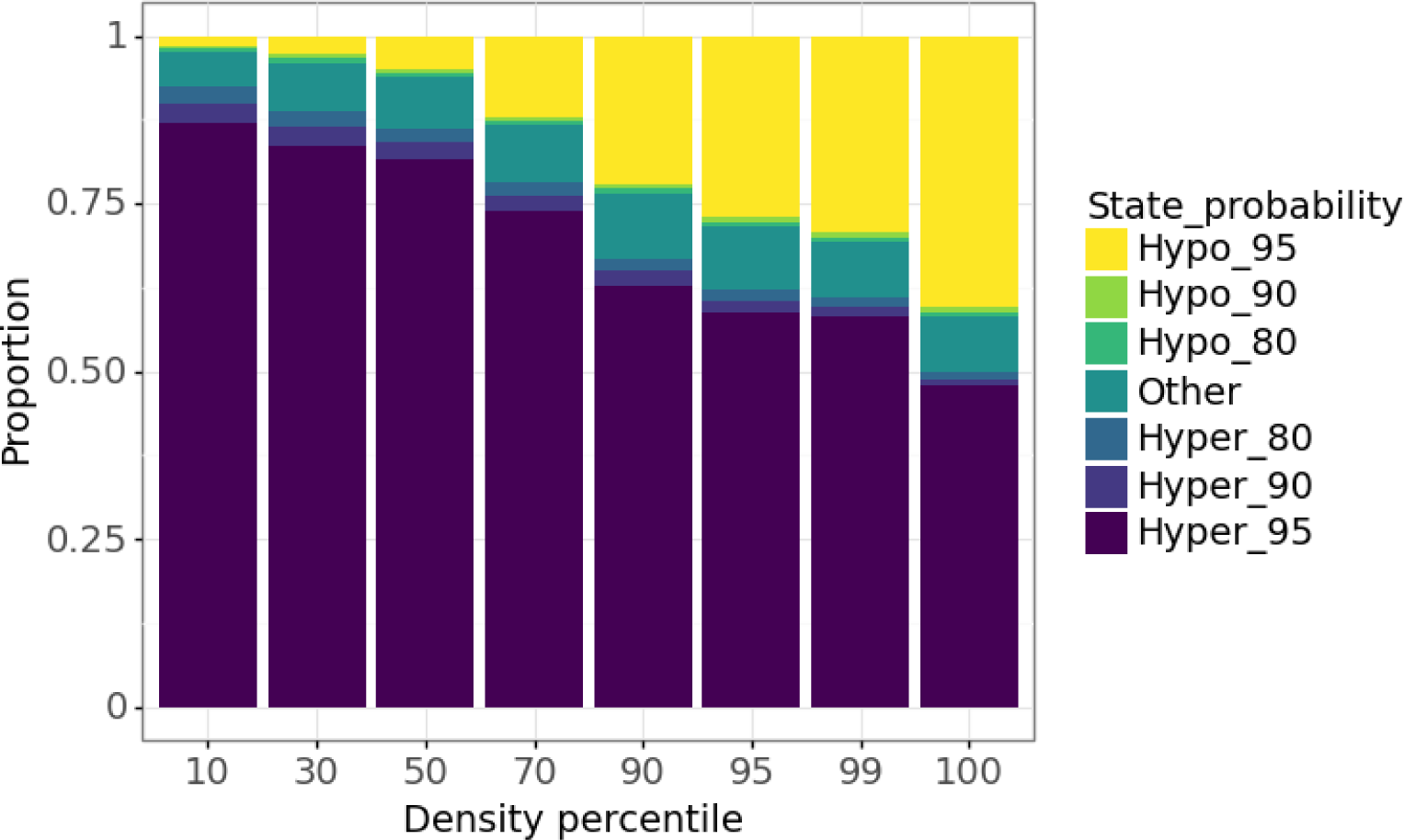
Distribution of inferred historical methylation status by local CpG density. We group all autosome CpGs into percentiles based on CpG density within 10kb (the leftmost bin summarizes 10% CpG sites that have the lowest CpG density neighborhood). Y-axis partitions CpGs in each density bin by their posterior probability of being hyper- or hypo-methylated, from having >0.95 probability of being in hyper-methylated state to having >0.95 probability of being in hypo-methylated state. About 10% CpGs are estimated to have intermediate methylation levels regardless of local densities.

**Figure A.2:**
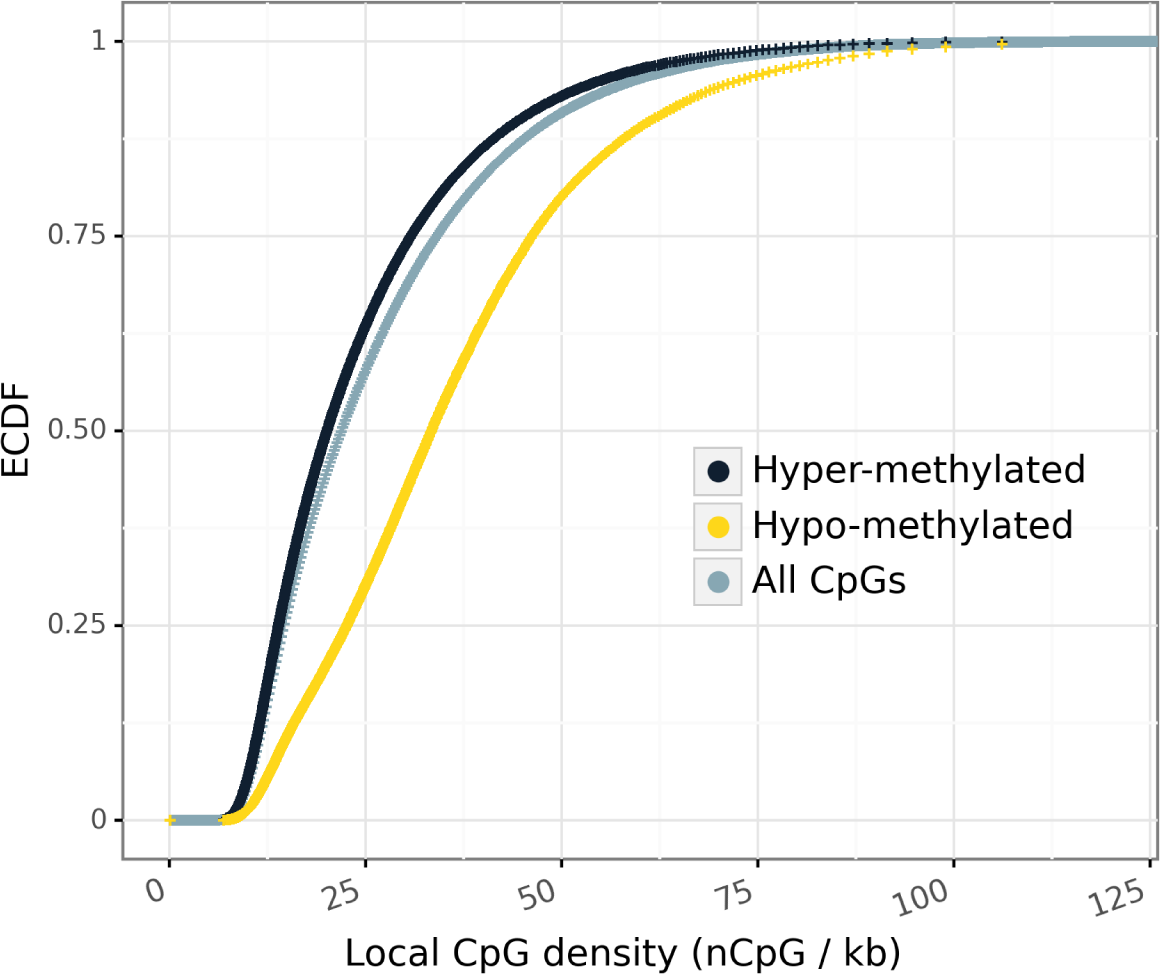
Distribution of local CpG densities measured as the number of CpG sites within *±*10kb of a focal CpG. Methylation status is based on MHMM. Hypo-methylated CpGs tend to locate in denser regions.

**Figure A.3:**
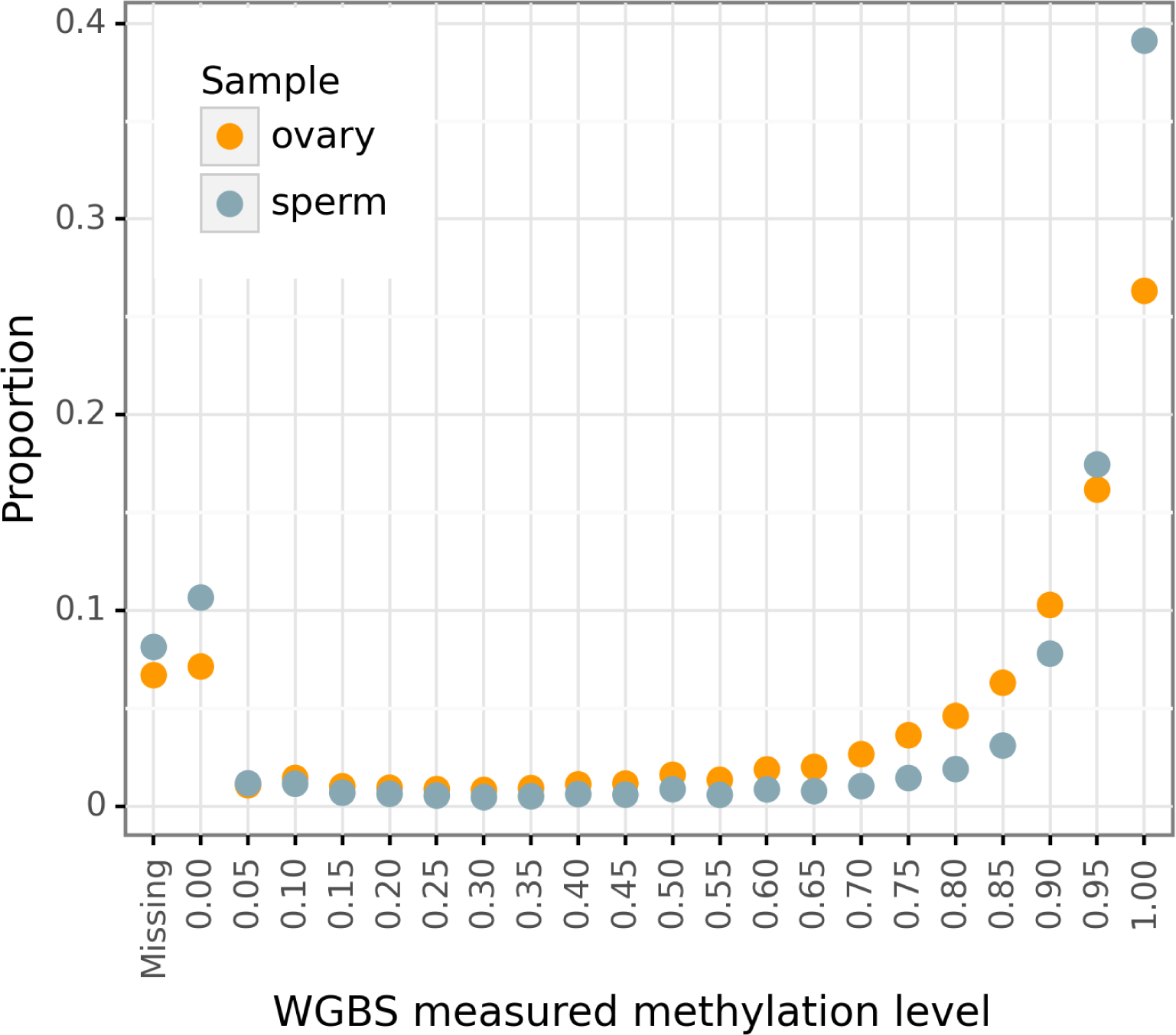
Distribution of methylation status measured in two high quality WGBS germline samples. X-axis indicates the upper bound of 0.05 intervals (right closed), leftmost point indicates the missing rate. Proportion is among 55.3M autosome CpGs.

**Figure A.4:**
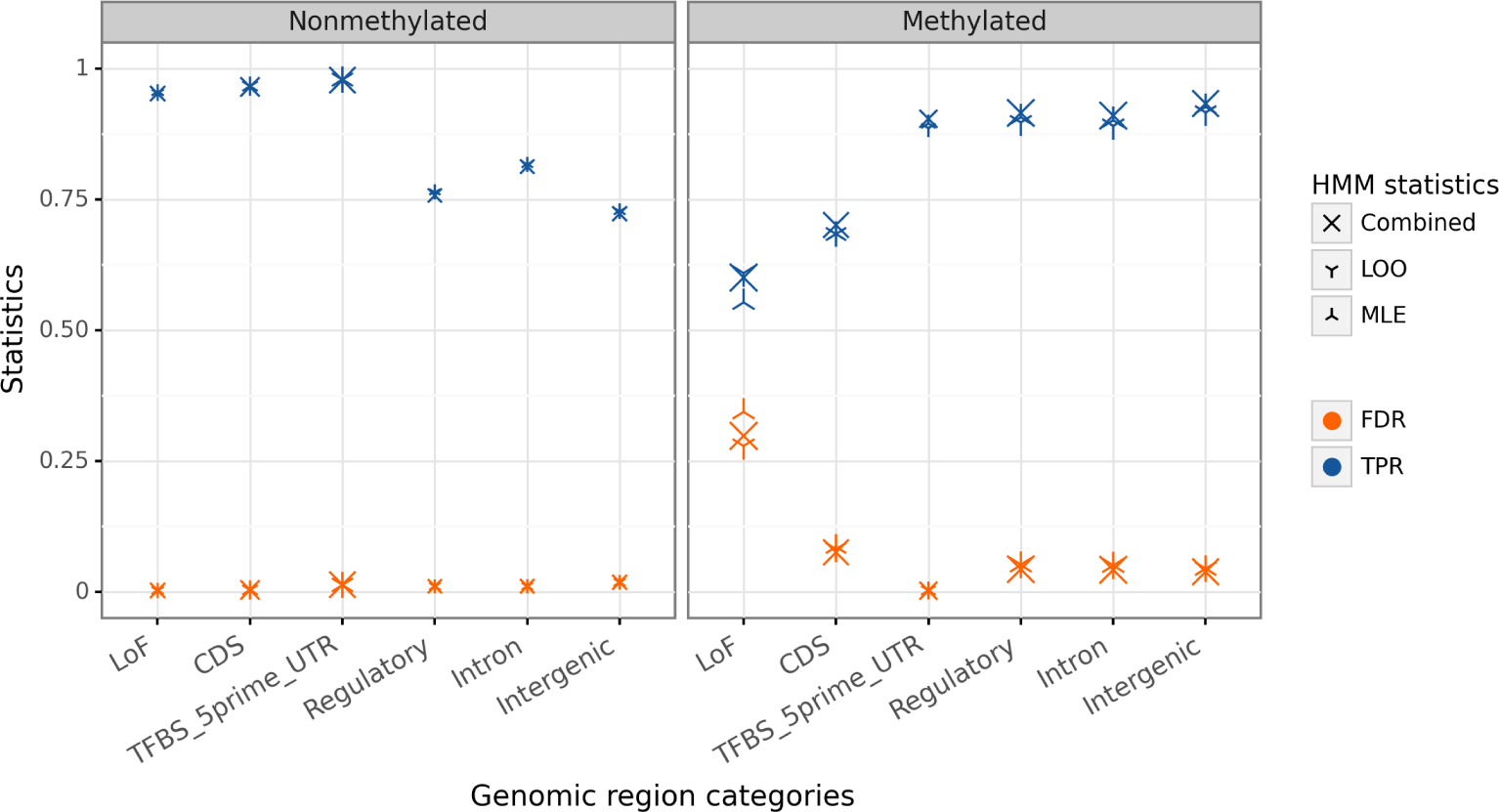
Comparing HMM estimates with WGBS measures stratified by functional regions. Here we treat WGBS as the “true” label to define TPR and FDR for comparison. The two methods differ more in non-active hypo-methylated sites and negatively selected sites. (TPR: MHMM inferred hyper-methylated CpGs among WGBS identified methylated sites. FDR: WGBS identified non-methylated sites among MHMM inferred hyper-methylated CpGs.)

**Figure A.5:**
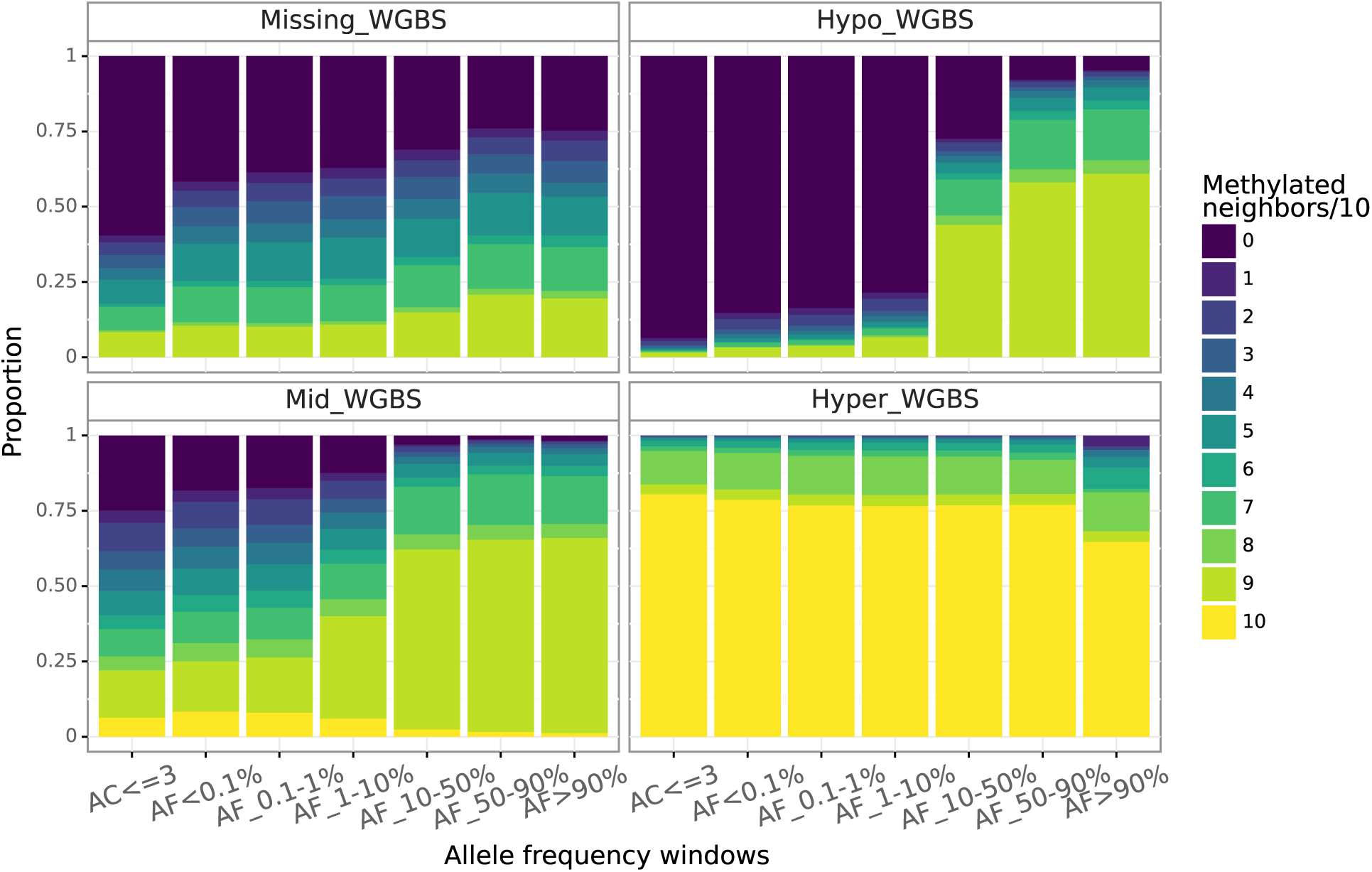
Methylation pattern of the close neighborhood around any focal CpG sites measured by WGBS in a sperm cell line. We stratify the focal CpG according to its measured methylation level and allele frequency in Bravo (x-axis), then within each stratum visualizes the distribution of the number of methylated neighbors among *±*5 nearby CpG sites. A focal site methylation status is in general consistent with its neighbors except for high allele frequency non-methylated CpGs, which are observed much more often in methylated neighborhoods. (WGBS has 2bp resolution, so a non-methylated site has at most 9 out of its 10 neighbors methylated)

**Figure A.6:**
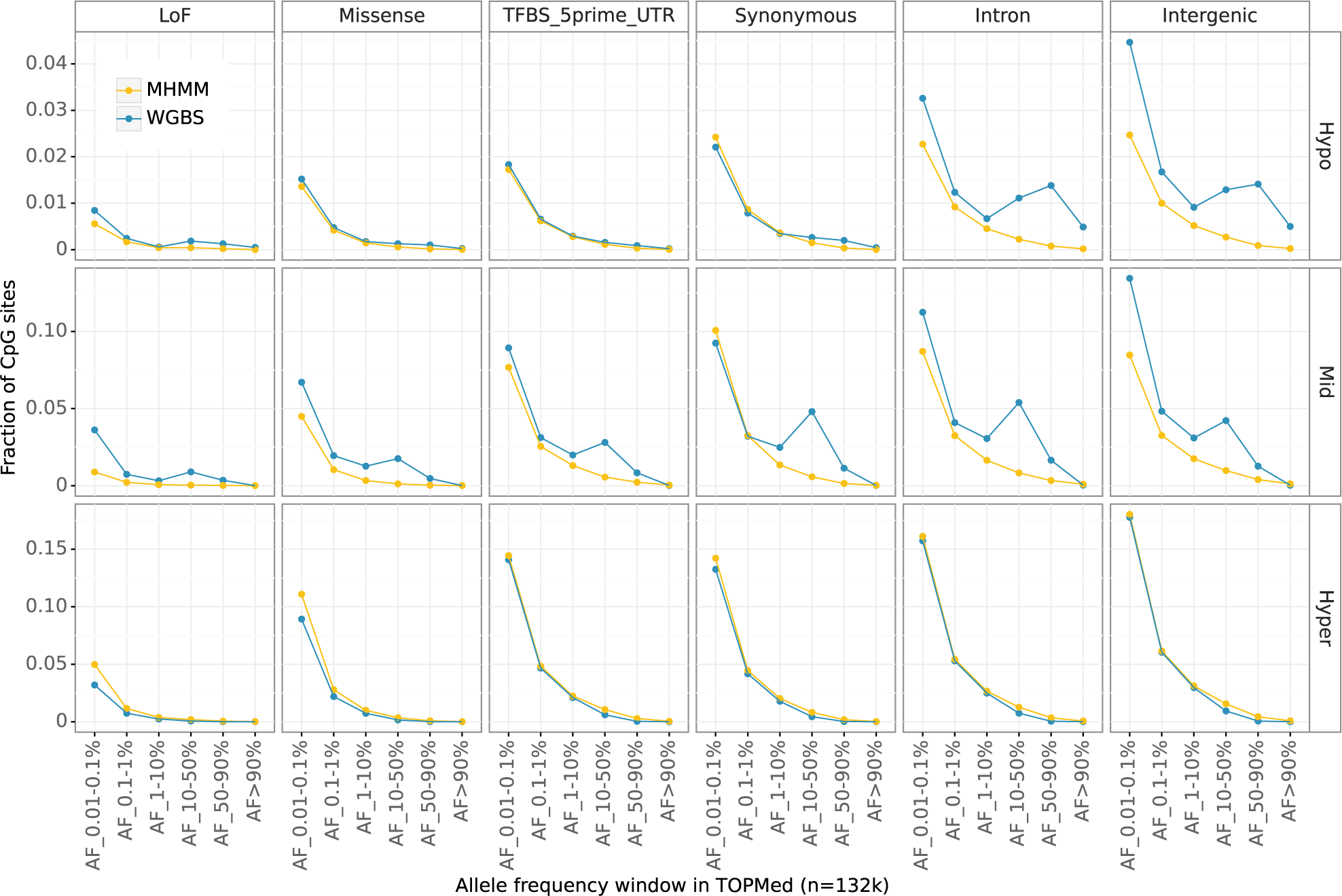
SFS of CpG C>T mutations stratified by MHMM inferred germline (yellow) or WGBS measured sperm (blue) methylation levels (rows) and predicted variant effects from Ensembl VEP[30] (columns). Y-axis shows the fraction among all CpG sites in the reference genome; fractions of sites with AF < 0.1% are not shown to focus on the difference in intermediate and high frequency variants.

**Figure A.7:**
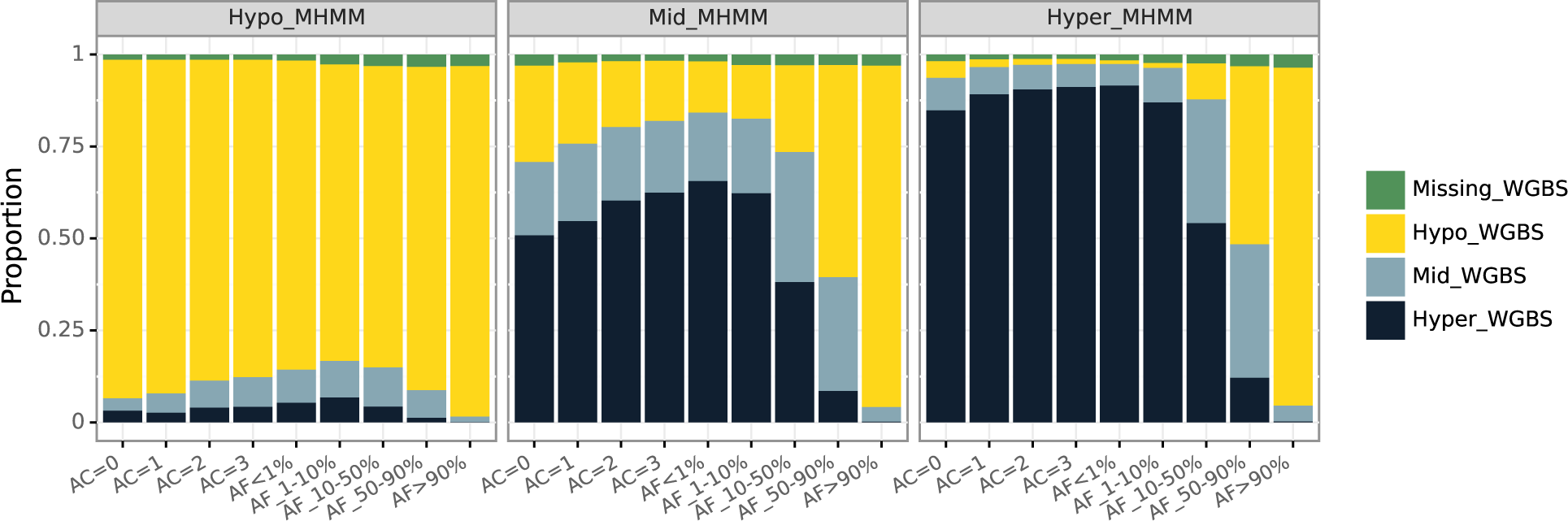
Composition of WGBS measured methylation status among CpGs in each MHMM inferred state (three panels) and allele frequency categories (x-axis). X-axis shows the allele count (AC) or allele frequency (AF) of the T allele at CpG sites (data from Bravo), from monomorphic (left most) to high frequency (right). The high discordance between inferred methylation status and measured methylation status for high T allele frequency illustrates the germline mutation bias of WGBS.

**Figure A.8:**
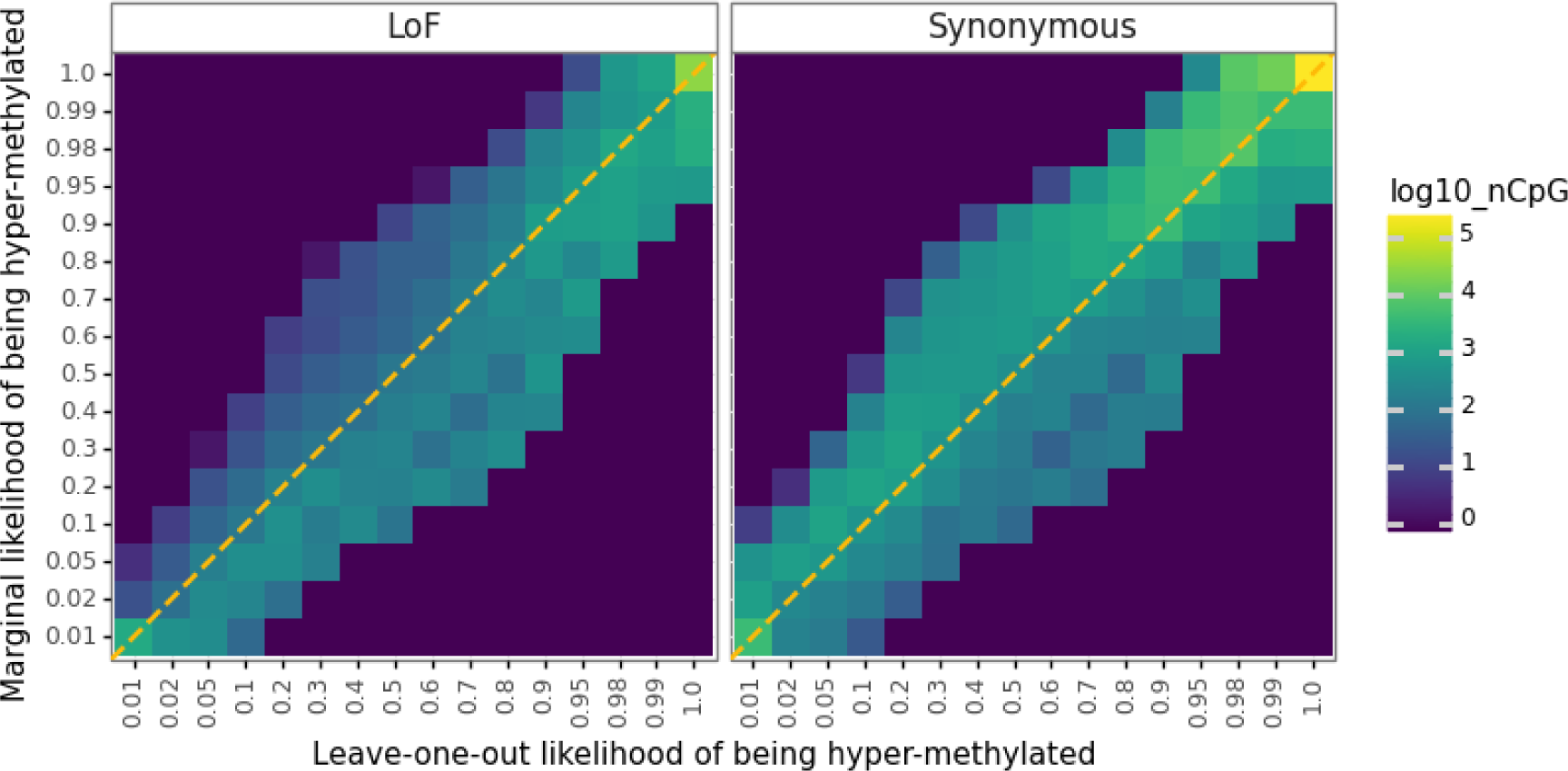
Joint density distribution of leave-one-out likelihood (x-axis) and the marginal likelihood (using in addition the focal site’s allele frequency, y-axis) of hyper-methylation at CpGs where the potential C>T mutations would be loss of function (LoF) or synonymous.Color indicates the number of CpGs in each discretized parameter window. At LoF variants, cells with large counts tend to be located below the diagonal, as the marginal likelihood infer hyper-methylation less confidently than leave-one-out likelihood. At synonymous variants, cells with large counts tend to be located above the diagonal, as the marginal likelihood infer hyper-methylation more confidently than leave-one-out likelihood.

**Table A.1:** Compare MHMM results from Bravo v.s. those from gnomAD. Numbers in the table are fractions among all analyzed autosome CpGs (45M). Fractions of CpGs that the results from two datasets agree are highlighted by green

| MHMM in gnomAD | MHMM Hyper in Bravo |  |  | MHMM Hypo in Bravo |  |  | MHMM Mid in Bravo |  |  |
| --- | --- | --- | --- | --- | --- | --- | --- | --- | --- |
|  | Hyper | Hypo | Mid | Hyper | Hypo | Mid | Hyper | Hypo | Mid |
| Hyper_WGBS | 0.7032 | 0.0002 | 0.0190 | 0.0034 | 0.0005 | 0.0007 | 0.0303 | 0.0010 | 0.0059 |
| Hypo_WGBS | 0.0148 | 0.0002 | 0.0020 | 0.0015 | 0.0998 | 0.0018 | 0.0059 | 0.0098 | 0.0054 |
| Mid_WGBS | 0.0548 | 0.0002 | 0.0029 | 0.0015 | 0.0029 | 0.0009 | 0.0083 | 0.0026 | 0.0049 |
| Missing_WGBS | 0.0120 | 0.0001 | 0.0003 | 0.0003 | 0.0009 | 0.0002 | 0.0011 | 0.0003 | 0.0004 |
| Column sum | 0.7848 | 0.0005 | 0.0242 | 0.0067 | 0.1041 | 0.0036 | 0.0457 | 0.0137 | 0.0166 |

**Table A.2:** Hypo- but not hyper-methylated CpGs are enriched in known active elements. Hypo and hyper labels are based on MHMM, indicating ≥ 90% CpGs in the CpG island are hypo- or hyper-methylated respectively. Numbers in the table are the proportion of CpG islands overlapping with known active elements.

| Feature | CpG islands |  |  |
| --- | --- | --- | --- |
|  | Fraction that overlaps with known features |  |  |
|  | Hypo (19842) | Other (2447) | Hyper (3454) |
| Promoter | 0.4221 | 0.0327 | 0.0035 |
| Proximal enhancer | 0.3700 | 0.0372 | 0.0055 |
| ATAC-seq peak (open chromatin) | 0.3488 | 0.0670 | 0.0217 |
| TF ChIP-seq (TF binding sites) | 0.1618 | 0.0327 | 0.0136 |
| Any of above | 0.6831 | 0.1300 | 0.0431 |

**Table A.3:** Purifying selection maintains CpGs as monomorphic at functional sites. Numbers in the table are fractions of monomorphic sites conditional on the CpG’s functional annotation of the potential T allele (where the C in a CpG site is the reference allele) and methylation level. The light blue cells highlight the fraction of monomorphic hyper-methylated CpG sites where the C>T mutation is predicted as loss-of-function, where the methylation levels are from MHMM (left) and sperm WGBS (right) respectively.

| Functional category | All CpGs (45.2M) | Monomorphic proportion |  |  |  |  |  |  |
| --- | --- | --- | --- | --- | --- | --- | --- | --- |
|  |  | MHMM inferred methylation level |  |  | Sperm WGBS methylation level |  |  |  |
|  |  | Hyper (34.9M) | Mid (4.8M) | Hypo (5.5M) | Hyper (34.5M) | Mid (3.6M) | Hypo (6.4M) | Missing (0.7M) |
| LoF (74550) | 0.4773 | 0.2794 | 0.6162 | 0.7810 | 0.4330 | 0.4541 | 0.7492 | 0.3528 |
| Missense (1337129) | 0.3155 | 0.1078 | 0.2878 | 0.6668 | 0.1737 | 0.2413 | 0.6632 | 0.3088 |
| TFBS or 5' UTR (806960) | 0.4962 | 0.0692 | 0.2196 | 0.6679 | 0.0744 | 0.1961 | 0.6607 | 0.3156 |
| Synonymous (583647) | 0.2612 | 0.0655 | 0.1223 | 0.6102 | 0.0769 | 0.1505 | 0.6229 | 0.2715 |
| Regulatory (2432178) | 0.1202 | 0.0603 | 0.1973 | 0.5559 | 0.0664 | 0.1543 | 0.4850 | 0.1694 |
| Intron (26595687) | 0.1287 | 0.0599 | 0.1972 | 0.6090 | 0.0682 | 0.1429 | 0.5341 | 0.1550 |
| Intergenic (13415783) | 0.1257 | 0.0512 | 0.2267 | 0.5917 | 0.0571 | 0.1268 | 0.4735 | 0.1573 |

**Table A.4:** (In Excel file) Overall methylation rates and comparison with MHMM results in 70 publicly available WGBS samples from ENCODE. Ages of the donors of the germline samples are listed in parentheses. The first two samples are the same as those used in [5]. We only include samples with missing rate less than 20% in autosomes (after removing CpGs with low mapping or sequencing qualities).

## B List of public materials

- gnomAD[6] v3.0 site lists and allele frequencies: https://gnomad.broadinstitute.org/
- Bravo[14] (TOPMed freeze 8) site lists and allele frequencies: https://bravo.sph.umich.edu/freeze8/h
- Germline CpG methylation from ENCODE[16] Sperm primary cell: ENCODE ID ENCSR705FPH (GEO accession GSM1127119) Ovary tissue: ENCODE ID ENCSR417YFD (GEO accession GSM1010980)
- Functional annotation from Ensembl VEP[30]: https://useast.ensembl.org/info/docs/tools/vep
- Transcription start sites from FANTOM5[37]: https://fantom.gsc.riken.jp/5/
- Open chromatin annotation from ENCODE[16]: ATAC-seq in testis (one male), identifier ENCSR210NKB TF CHiP-seq (same individual as the above ATAC-seq data), identifier ENCSR753RME
- WGBS datasets from ENCODE[16] see Table A.4

